# Dynamic selective auditory attention detection using RNN and reinforcement learning

**DOI:** 10.1101/2021.02.18.431748

**Authors:** Masoud Geravanchizadeh, Hossein Roushan

**Affiliations:** Faculty of Electrical & Computer Eng., University of Tabriz, Tabriz, Iran, 51666-15813

**Keywords:** selective auditory attention detection, dynamic learning system, recurrent neural networks, reinforcement learning, EEG, cocktail party problem

## Abstract

The cocktail party phenomenon describes the ability of the human brain to focus auditory attention on a particular stimulus while ignoring other acoustic events. Selective auditory attention detection (SAAD) is an important issue in the development of brain-computer interface systems and cocktail party processors. This paper proposes a new dynamic attention detection system to process the temporal evolution of the input signal. The proposed dynamic SAAD is modeled as a sequential decision-making problem, which is solved by recurrent neural network (RNN) and reinforcement learning methods of Q-learning and deep Q-learning. Among different dynamic learning approaches, the evaluation results show that the deep Q-learning approach with RNN as agent provides the highest classification accuracy (94.2%) with the least detection delay. The proposed SAAD system is advantageous, in the sense that the detection of attention is performed dynamically for the sequential inputs. Also, the system has the potential to be used in scenarios, where the attention of the listener might be switched in time in the presence of various acoustic events.

## 1. Introduction

For years, neurocognitive scientists have made great profits from segregating the brain into different functioning domains. The behavior of an organism aimed toward a task requires the joint operation of memory, executive functioning, attention, language, and sensorimotor units [1]. Attention is the foundation for all the other cognitive functions that deal with the ability to focus on distinct aspects of information or awareness on a given stimulus or task, long enough to accomplish a goal and to shift awareness, if appropriate. This means that the human listener can shift his attention both consciously and sometimes unconsciously in response to the environment. Auditory selective attention is the process in which a person attends to one or a few sounds while ignoring the other ones; a phenomenon called the cocktail party problem. The first formal description of the cocktail party problem was given by the psychologist Cherry in 1953 by demonstrating various dichotic experiments [2]. Cherry conducted attention experiments in which participants listened to two different messages from a single loudspeaker at the same time and tried to separate them; this was later termed dichotic listening task. It is believed that in a high-level auditory cognitive process, two interacting critical mechanisms are involved in the identification of sounds in a complex auditory scene. These include sound segregation, also called auditory scene analysis (ASA), and attentional selection [3]. According to the perceptual process of ASA, the sound mixture is decomposed into a collection of segments which are subsequently grouped to form coherent streams, a procedure known as object formation [4–6]. The studies show that attention operates on auditory objects and the desired object is selected by the direction of top-down attention [7, 8]. Nevertheless, there is as yet little understanding as to the role of auditory scene analysis and auditory attention in the identification of sound and the argument about the relation of object formation and object selection is ongoing [3].

The understanding of neurobiological solutions of the cocktail party problem by the brain, and also the recent technological advances make it possible to explore potential applications of selective auditory attention detection (SAAD). A few examples of practical applications include enhancing the performance of speech separation algorithms, cognitive hearing aids, brain-computer interface (BCI) systems, etc. among others.

There are many reports that auditory attention can be detected from brain signals, using various neural signal acquisition, including non-invasive magnetoencephalography (MEG) [9], electroencephalography (EEG) [10], and invasive electrocorticography (ECoG) [11, 12]. EEG signals can be considered as the reflection of electrical activity in the cerebral cortex which contains a wealth of information related to advanced nervous activities in the human brain such as learning, memory, and attention [13]. The advantages of relatively low cost, easy access, and high temporal resolution make EEG signal acquisition a valuable candidate for the study of auditory attention [14].

Much research has been conducted to study SAAD based on EEG data which is produced from non-speech stimuli. In this context, the effects of attention on auditory sound segregation have been investigated using event-related potentials (ERPs). These studies show that selective attention may operate in a two-stage process, including an early stage of bottom-up stream separation based on acoustical features of sounds, and a later top-down task-dependent stage [15–17]. SAAD generated from non-speech stimuli has been also the focus of studies by some researchers in the framework of auditory steady-state response (ASSR). ASSR is a brain activity response typically obtained by periodic amplitude modulated sinusoidal tones or click sound trains as auditory stimuli [18–20]. The major disadvantage of the ERP and ASSR methods is that they are unsuccessful for natural continuous speech stimuli [21–23] which occurs in real environments. The human auditory system has efficiently evolved to attentively focus on a salient stimulus of an auditory scene occurring in cocktail party scenarios where most of the sound sources are continuous natural speech.

As one method of decoding the attentional direction employing natural speech, some researchers have conducted SAAD experiments by machine learning techniques using the extracted informative features of EEG signals for training classifiers [7, 14, 22, 24]. As yet, different informative features have been employed in the design of classifiers. Recently, the benefits of connectivity measures for the detection of selective auditory attention were introduced by extracting optimized features based on the Granger causality approach [25]. The main advantage of this method is that the classification of the attentional state is performed from single-trial EEG signals without reconstructing the speech stimuli. Stimulus-response modeling using temporal response functions (TRFs) has made important contributions in decoding the auditory attention of a listener in a competing-speaker environment. TRFs could be estimated by system identification approaches to quantify the mapping between amplitude envelopes of speech and EEG [26–28]. Some existing TRF-based techniques attempt to track the attentional state of a listener in a complex auditory environment by reconstructing attended and unattended speech in the low-frequency range (1 □ 8 Hz) using high-density EEGs [10, 29, 30]. In this frequency range, EEG corresponds to the spectrum of speech envelope. Here, the subject’s attention is detected based on the correlation between the reconstructed speech envelope and the actual attended and unattended speech envelopes at the two ears. In a similar study, using limited training data, Miran *et al.* [26] developed an algorithm for detecting the attentional state which consists of estimating real-time encoding/decoding coefficients, extracting attentional state markers, and implementing a near real-time state-space estimator. Alternatively, the concept of TRFs has been previously employed in the analysis of the human auditory system to describe the properties of such a system using EEGs [28, 31, 32]. Here, in a mapping process from speech features to neural data, TRFs could be used to predict EEGs from the attended speech envelopes. Power *et al*. [32] proposed the technique of auditory evoked spread spectrum analysis which extracts high-resolution temporal responses of two simultaneously presented speech streams in a condition most similar to a natural cocktail party environment. Recently, studies on the use of neural networks in SAAD have introduced new frameworks for decoding the listener’s attention [33, 34]. De Taillez *et al.* investigated non-linear machine learning methods such as deep neural network (DNN) with a novel architecture to replace the linear regression used in previous studies (e.g., O’sullivan *et al*., 2014) with the aim of better decoding of listener’s attention. In [34], inspired by the work of de Taillez *et al.*, a convolutional neural network (CNN) is used in the classification architecture. In this research, a different end-to-end decision network is used as the attention decoder with integrated similarity computation between EEG signals and a candidate audio envelope. In this paper, a novel dynamic SAAD is addressed to model the attention detection as a sequential decision-making problem with the involvement of time to process sequences of inputs, where dynamic learning methods are employed to describe the temporal evolution of the system. Here, a methodology is presented to answer the following questions:

a. Compared with non-dynamic approaches, to what extent are dynamic learning methods effective in the analysis of sequential data for the SAAD task?
b. Does the strategy of trial and error used in agent-based dynamic methods improve the ability of attention detection for having intelligent learning machines?
c. How do such dynamic systems perform in examining the attentional direction of listeners when their focus on speech stimuli is switching over time?

In this regard, the dynamic learning approaches of the recurrent neural network (RNN) and reinforcement learning (RL) are incorporated in the detection model. RNN is used to process input sequences and can be termed as a DNN in the “temporal” sense [35] to make direct decisions of which speech stream the listener is focused on at any moment. In RNN, unlike feedforward neural networks, the outputs of each layer are fed back into the inputs of previous layers, which provides the characteristics of a system with memory [36]. As a second dynamic learning approach, the concept of RL is employed in the attention detection process, formalized by the Markov decision process (MDP) framework and solved by *Q*-Learning (QL) and deep *Q*-learning (DQL). The RL-based system is composed of a set of agents that learn to create successful strategies using rewards in a trial and error procedure [37]. In an inspiring study [38], the problem of classifying imbalanced data is modeled as a sequential decision-making problem which is solved by DQL. Instead of the traditional classification process in which extracted features from the input are used to estimate the class label, here, an agent is used to interact with the environment. The use of dynamic system architecture in the attention detection task yields the flexibility to investigate the attention-switching behavior of listeners and introduces a framework for tracking attention in real-time applications while the focus of the listener is shifting between streams over time. The organization of the paper is as follows. Section 2 explains the methodology, including the data description and the proposed dynamic selective auditory attention detection model. Here, the details of the model, including the structure of probabilistic state-space, and the learning methods are described. In Section 3, the experiments and evaluations along with discussions are given. The concluding remarks and some perspectives for future work are presented in Section 4.

## 2. Materials and methods

### 2.1 Data description

In this work, the publicly available data of 20 normal-hearing subjects are used for the evaluation of the experiments [39]. Two different stories are presented simultaneously via a headphone to each subject: one to the left ear and the other to the right ear, and the subjects are asked to attend aurally to just one of the stories, where at the same time the EEG of participants with 128 electrodes are recorded. In each trial, half of the subjects are asked to attend to the speech on the left ear and the remaining half to the speech on the right ear. Three trials are considered for each person, each having a duration of approx. 60 seconds. To ensure that the subjects have performed their attentional tasks correctly, a questionnaire about the information of the stories is used. The EEG signals are recorded at 128 Hz sampling rate, average referenced, and finally preprocessed to minimize the presence of 50 Hz line noise, eye blink, and muscle movement artifacts.

### 2.2 Proposed SAAD system

The proposed dynamic selective auditory attention detection model is shown in Figure 1. Here, after the preprocessing of the input signals, the probabilistic state space which is the set of all possible states of the system is formed. The values of the state variables at a particular time gives the state of the system at that time. Next, in the learning stage, based on the computed state variables and the available true labels of the attention directions, three different machine learning methods are applied to make the final decision as to the attended speaker. Due to the dynamic nature of the learning methods for the proposed SAAD model, there is no training stage for the classifier.

**Figure 1:**
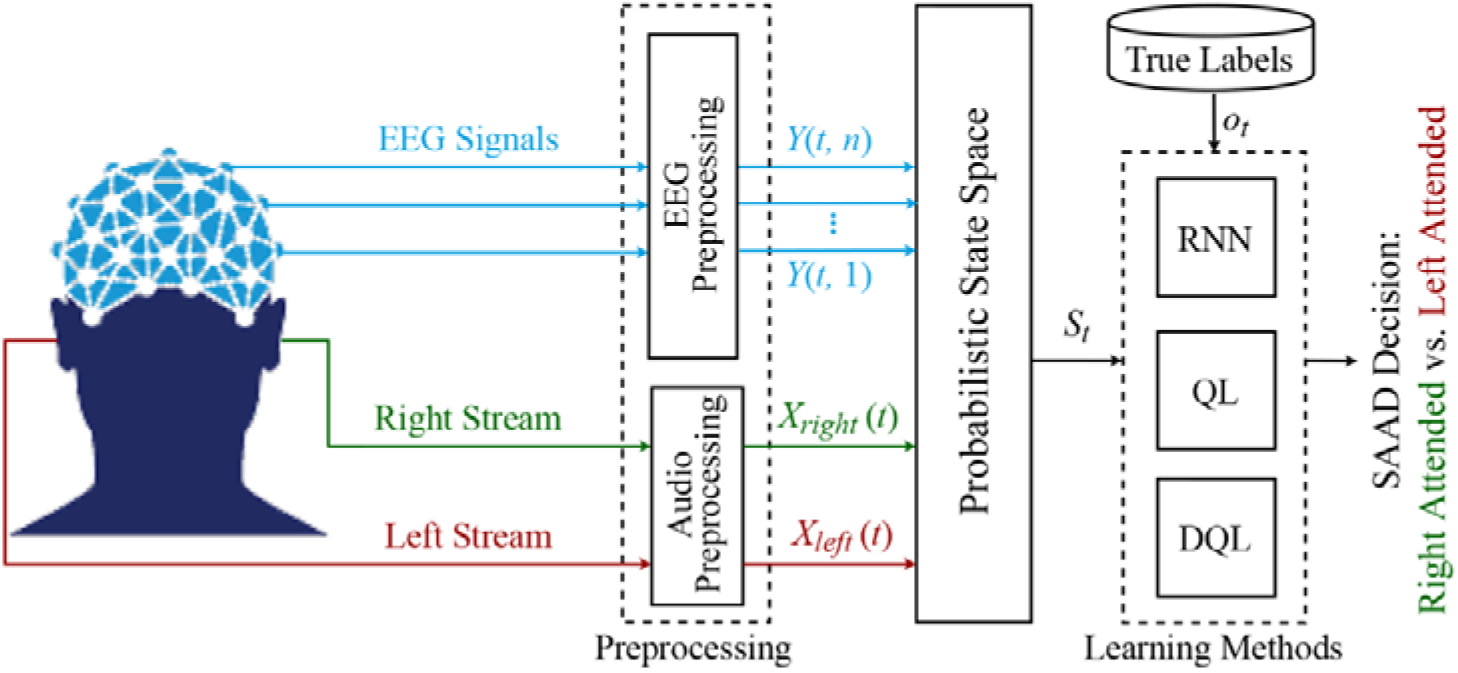
The block diagram of the proposed SAAD system using the dynamic learning methods of RNN, QL, and DQL.

#### 2.2.1 Preprocessing

In this stage, EEG signals are filtered with band-pass filters in the range of 2 □ 30 Hz to obtain the useful cognitive information of EEG data in this frequency range. Then, both EEGs and speech stimuli are downsampled to the same sampling rate of 64 Hz to decrease the required processing time of later stages.

#### 2.2.2 Probabilistic state space

One of the main goals of computational neuroscience is to develop techniques to characterize the dynamic features inherent in cognitive tasks that have rich temporal structures. A complete description of a dynamical learning system can be given by a set of variables whose values at a particular time yields the state of the system at that time. These variables define the state of the system and the set of all their possible values is called the state space of the dynamical system [40]. State spaces are highly descriptive for learning patterns in time series data. The state-space representation gives a suitable and compact way to model and analyze systems with multiple inputs and outputs. Probabilistic state-space models (P-SSMs) provide a general framework for analyzing stochastic dynamical systems that are observed through a stochastic process. P-SSMs describe systems at the time *t* with input **X**_*t*_ and output **Y**_*t*_ in terms of a Markovian state *S*_*t*_, based on an observation model *f* and transition model *g* [41]:

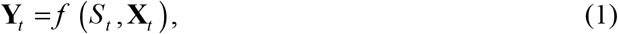

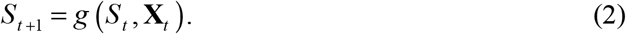

Here, **X**_*t*_ = [*X*_*left*_ *(t)*, *X*_*right*_*(t)*] represents the left- and right-ear stimuli and **Y**_*t*_ = [*Y* (*t*, 1),….,*Y (t, n)*] is the EEG responses measured for *n* electrodes (*n* = 128). The computed Markovian state *S*_*t*_ is fed to the learning methods of the successor stage.

#### 2.2.3 Learning methods

##### 2.2.3.1 RNN

A recurrent neural network is a multi-layer neural network used to analyze sequential input for classification and prediction purposes. Different from feedforward neural networks, RNNs are not limited by the length of input and can use their internal state (i.e., memory) to process sequences of inputs. The RNN architecture considers the current input and the output learned from the previous input to make a decision.

The general structure of the RNN learning system is depicted in Figure 2. As shown, the learning system takes some input state *S* at a particular time *t* and feeds that input into the hidden layers having an internal state, *h*, at that time. The values of the hidden states are fed back to the learning model and updated every time RNN receives a new input. At each time step, the current hidden state, *h*_*t*_, is updated by the previous state, *h*_*t*–1_, and the current input, *S*_*t*_, based on the recurrence formula [42]:

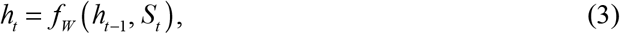

where
*f*_*W*_ is defined typically by “tanh”, as the activation function, and a set of weights, *W*:

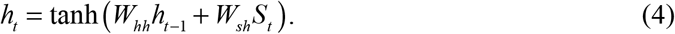

**Figure 2:**
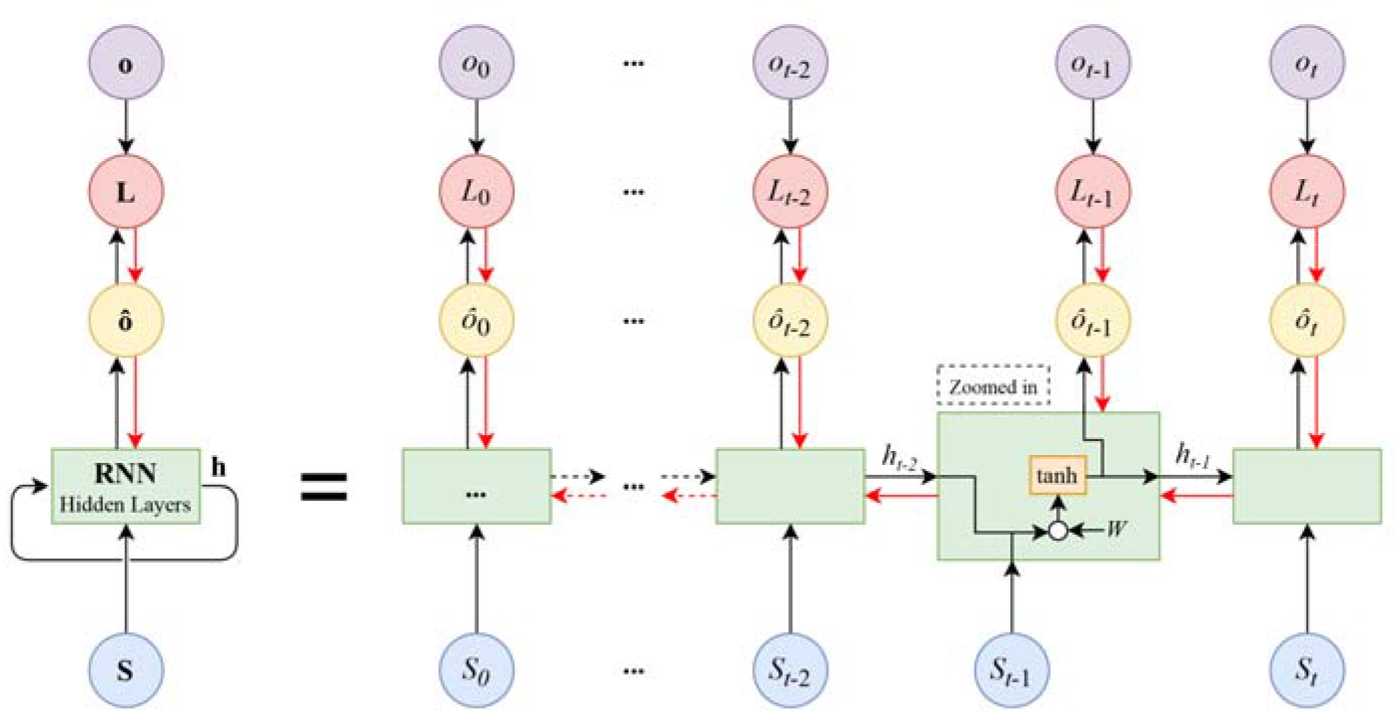
The block diagram of the RNN learning system is shown in rolled (left) and unrolled (right) configurations. The black arrows illustrate the forward propagation path, whereas the red arrows depict the backpropagation path. A zoomed view of a sample hidden layer is shown to display the detailed internal structure.

**Figure 3:**
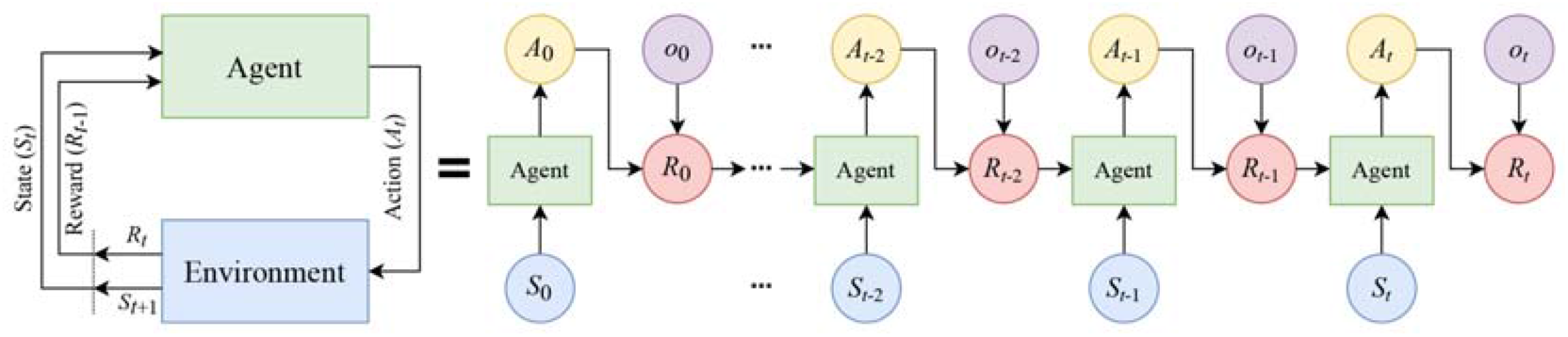
The block diagram of MDP in rolled (left) and unrolled (right) configurations. The agent and the environment interact over a sequence of discrete-time steps. At each time step, the agent receives a representation of the state *S*_*t*_ of the environment and a reward *R*_*t*–1_ from the previous interaction to issue an action *A*_*t*_.

**Figure 4:**
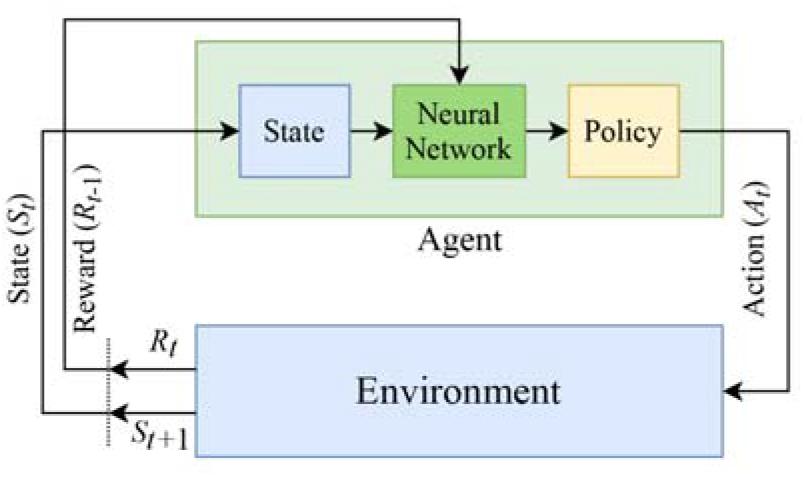
The block diagram of the DQL system. The key difference with QL is the use of a neural network in the learning process of the agent.

The predictions of output *ô* as the objective of RNN at each time step are computed as:

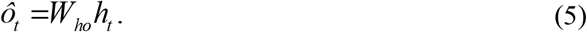

The parameters *W*_*hh*_, *W*_*sh*_, and *W*_*ho*_ are the weights of the RNN architecture which are shared throughout the entire network and initialized with random values. RNN uses the backpropagation method to learn from sequential training data to correct its prediction. This is achieved by updating the weights using the gradients of a computed loss function, *L*, between prediction, *ô*, and true value, *o*. The learning procedure is continued until the loss value is reduced to a certain threshold, called the stop threshold, after which the backpropagation process stops.

Referring to Figure 1 and using the learning method of RNN as the classifier in the proposed dynamic SAAD system, at each time step *t*, the output of probabilistic state space, *S*_*t*_, is given to the hidden layers of RNN. At this time step, the RNN classifier predicts the class label, *ô*_*t*_, of the attentional direction. The predicted value is compared with the true attentional labels, *o*_*t*_, based on a predefined loss function. The internal weights of RNN are updated through the backpropagation process as long as the loss function is higher than a certain threshold before the state at the next time step is fed into RNN. The values of *ô*_*t*_ at the output of RNN are the predicted labels specifying the left or right attended speech. The parameters characterizing the detailed structure of the RNN learning method are shown in **Error! Reference source not found.** (Sec. 3.1).

##### 2.2.3.2 Reinforcement learning

Reinforcement learning is a dynamic learning method dealing with the design of intelligent agents that learn through trial and error strategy by interacting with their environment. The general operation of RL is based on a sequence of states, actions, and rewards. The typical structure of RL consists of an environment that represents the outside world and an agent that takes actions based on received observations from the environment. The environment includes the current state and a reward that informs the agent of how good or bad was the previous action to improve its performance. The RL task can be formalized as Markov decision process and solved by *Q*-Learning and deep *Q*-learning which are described in detail below.

###### A. Q-Learning

Almost all RL problems can be formalized as a Markov decision process which is a discrete-time state-transition system. In MDP, the environment is stochastic and satisfies what is known as the Markov property. The Markov property states that given the current state and action, the next state is independent of all previous states and actions. MDPs can be described formally with the following components: S denotes the state space of the process; A is the set of actions; P is the Markovian transition model, where *P*(*S*_*t*+1_ │ *S*_*t*_, *A*_*t*_) is the probability of making a transition to state *S*_*t*+1_ when taking action *A*_*t*_ in the state *S*_*t*_; R represents the reward function or feedback, *R*_*t*_, from the environment by which the success or failure of an agent’s actions is measured [43]. In MDPs, the behavior of the model is defined by the reward function. **Error! Reference source not found.** depicts the interaction between the agent and the environment in an MDP.

In QL, it is typical to compute a policy as the solution to the Markov decision process. A policy is a mapping from state to action (i.e., π: S → A). It indicates the action *A*_*t*_ to be taken while in the state *S*_*t*_. In the simplest case, the objective of RL is to find a policy that maximizes the discounted return *G*_*t*_ for each state which is the total discount reward from time-step *t* [43]:

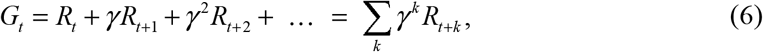

where 0 ≤ *γ* < 1 is the discount rate to balance the immediate and future rewards. Given that the discounted return function is stochastic, the expected discounted return, starting from state *S*, taking action *A*, and following policy *π*, is given as [44]:

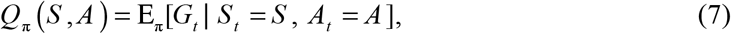

where *Q*_*π*_(*S*, *A*) is called the “action-value function” and denotes the expectation operator. The *Q*-value can be learned from a trial and error procedure, in which the agent may need to sacrifice small immediate rewards in exchange for the larger long-term ones. To this aim, the action-value function can be written in the form of a Bellman expectation equation. Using the Bellman equation, the function is decomposed into the immediate reward, *R*_*t*_, and the discounted *Q*-value of the successor state, *γ Q*_*π*_(*S*_*t*+1_, *A*_*t*+1_):

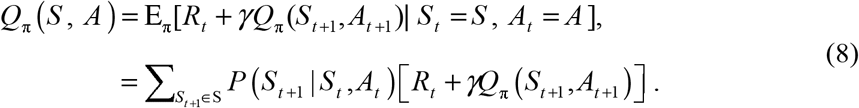

This equation expresses a relationship between the value of a state and the values of its successor states. The Bellman equation (Eq. (8)) establishes an iterative approach to calculate the optimal policy. The optimal policy *π*\* is the policy for which *Q*_*π*\*_(*S*, *A*) > *Q*_*π*_(*S*, *A*) among all possible policies *π*:

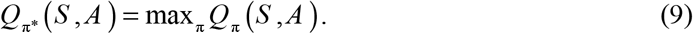

The function *Q*_*π*\*_(*S*, *A*) can be used to derive *π*\*(*S*):

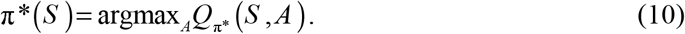

The optimal *Q*_*π*\*_ function can be found by inserting Eq. (9) into Eq. (8) [45]:

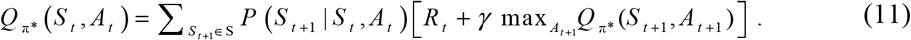

To estimate the action-value function, the method of *value iteration* can be adopted where the Bellman equation is updated iteratively:

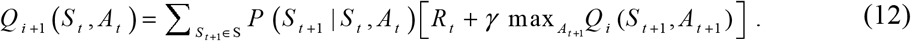

This algorithm converges to the optimal action-value function *Q*_π_ as the number of iterations increases (i.e., *i* → ∞).

###### B. Deep Q-Learning

The iterative Bellman equation (Eq. (12)) underlies many RL algorithms to estimate the action-value function. In practice, this basic approach may lead to instability, because the action-value function is estimated for each time sequence separately in which the samples would be highly correlated [46]. DQL can be regarded as an extension of the classical QL to approximate the optimal action-value function (i.e., *Q*_*π*\*_). In DQL, a history of interactions with the environment is used by the agent to learn the optimal policy. This type of RL algorithm employs a neural network as a function approximator (e.g., DNN), with weights parameter *θ*, called *Q*-network. The general block diagram of the deep *Q*-learning system is shown in **Error! Reference source not found.**.

The fundamental approach to solve the problem of instability in *Q*-networks is to break the temporal dependency and correlation among the sequence of observations used in training the neural network, called *experience replay* [47]. With experience replay, the agent’s experiences at each time step *t*, i.e., *e*_*t*_ = (*S*_*t*_, *A*_*t*_, *R*_*t*_, *S*_*t*+1_), are stored in a data set, called the replay memory. A *Q*-network can be trained by minimizing a loss function *L*_*i*_(*θ*_*i*_) defined as [46]:

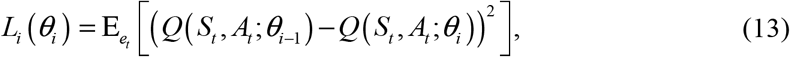

where *i* represents the iteration index. Using the gradient-descent approach, the optimal action-value *Q*-function is obtained when the minimum threshold value of the loss function is reached.

The two implementations of RL (i.e., QL and DQL) are used as the classifiers in the proposed dynamic SAAD system shown in Figure 1. Here, the output of the probabilistic state space, *S*_*t*_, and the true attentional labels, *o*_*t*_, form the environment. The classification agent uses the policy to predict the labels of the attention class represented by the action *A*_*t*_. At each time step, the agent receives a sample from the probabilistic state-space and classifies it. Then, the environment returns the next sample and an immediate reward *R*_*t*_ based on a comparison between the true and predicted classification labels. If the agent performs a correct classification action, which is the true detection of attention direction, it earns a positive reward (+1), otherwise, it is given a negative reward (−1). The agent’s task is to maximize the cumulative reward by learning optimal actions (i.e., true classification). This may involve sacrificing some initial immediate rewards to gain more long-term return. The policy of the classification agent is optimized by the Bellman iterative update for QL or minimizing the loss of the neural network (i.e., DNN or RNN in this study) for DQL. In Section 3.1, the detailed structures of both learning methods are shown in **Error! Reference source not found.**.

## 3. Experiments and evaluations

### 3.1 Experimental setup

To investigate the practical performance of the proposed dynamic SAAD model, in this study, three groups of experiments are implemented as follows. First, the efficiencies of different dynamic learning methods employed in the proposed model are evaluated. The internal structures of the learning methods used in the implementation of the proposed SAAD are shown in **Error! Reference source not found.**.

In the second group of experiments, the recently developed systems of attention detection in the literature [10, 27, 33, 34] are simulated as baselines and compared with the proposed SAAD system. The baseline systems are denoted, respectively, as “O’Sullivan *et al*.” [10], “Wong *et al*.” [27], “Taillez *et al*.” [33], and “Ciccarelli *et al.*” [34]. Although all the baseline systems use both EEG and speech signals as input, they have inherently different structures in the detection of attended speech. The method of “O’Sullivan *et al*.” uses a backward mapping technique to reconstruct the envelope of the attended speech. “Wong *et al*.” uses various TRF techniques to find a good regression in both forward and backward mapping. The baseline “Taillez *et al*.” employs DNN for the TRF regression, and “Ciccarelli et al.” uses convolutional neural networks (CNNs) for the learning of the end-to-end classifier. The proposed and baseline methods are simulated using the same EEG data obtained from 128 electrodes. Thirty percent of EEG data is used in the training procedure of the baseline systems and seventy percent of data is used to test them.

The last experiment concerns the applicability of the SAAD system in conditions where the attentional direction of the listener can be switched from one input stimuli to the other. To this aim, four artificial data sequences of 5, 10, 15, and 30 seconds with alternating attentional directions of the same subject (i.e., left-attended or right-attended) are created and concatenated to generate a whole sequence of 60 seconds long and used as input to investigate the performance of the proposed system in such switching attention conditions.

### 3.2 Performance measures

#### 3.2.1 Accuracy

In this study, the objective measure of accuracy is employed to validate the performance of classification. Accuracy (ACC) is a measure of the rate of total samples correctly classified by the model and is calculated as [48]:

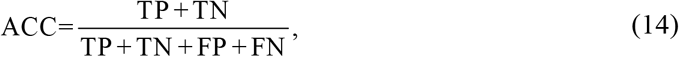

where true positive (TP) is the number of positive samples correctly predicted and true negative (TN) is the number of negative samples correctly predicted. False positive (FP) is denoted as the number of positive samples incorrectly predicted and false negative (FN) is denoted as the number of negative samples incorrectly predicted.

#### 3.2.2 Permutation test

In the classification studies, it is essential to reassure that the results are reliable in the sense that high detection performance is not due to overfitting. As an initiative in the reduction of the effects of overfitting, in this work, some basic methods such as multiple iterations of the algorithm and cross-validation techniques are employed. Yet, this does not necessarily mean overfitting has not occurred. Several studies suggest an evaluation approach, named as “permutation test” to confirm the competence of a classifier and validity of the results [49, 50]. In this approach, the attentional labels of the data (i.e., right attended/left attended) are permuted randomly to show that the whole classification pipeline fails with the new manipulated data. The reasoning behind the permutation test is to obtain accuracies with normal distributions centered on chance (i.e., 50% in 2-class problem) in multiple cross folding repetitions with the relabeled data [51]. The chance level accuracy of classification with randomly permuted labels illustrates that overfitting has not occurred in the detection of classes with the original data.

#### 3.3 Evaluations and discussion

In the first experiment, the performances of different learning methods are assessed in the detection of the attentional direction of the listener (see Figure 1). The results of the various classification approaches in terms of detection delays and ACC are shown in Table 2 obtained for 60-second trials and 100 repetitions of the proposed SAAD algorithm.

**Table 1:**
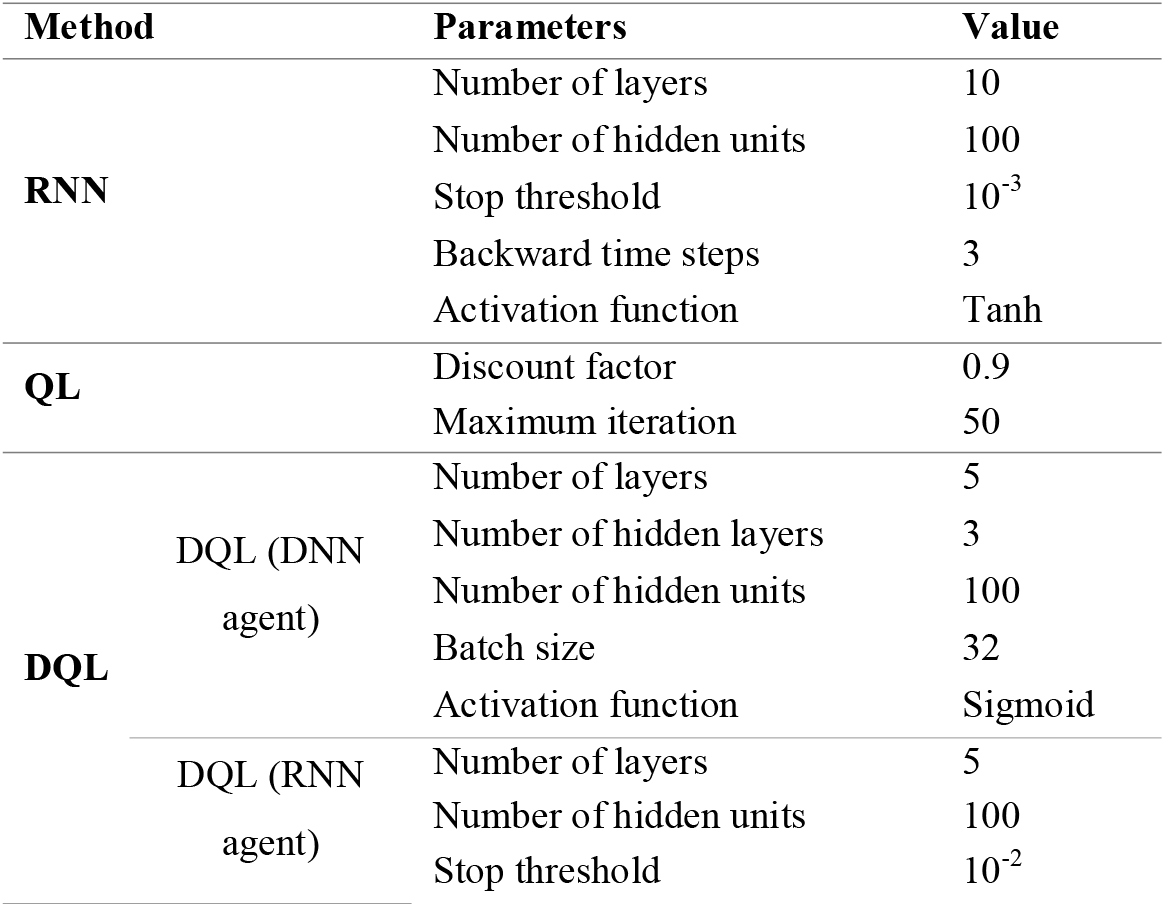
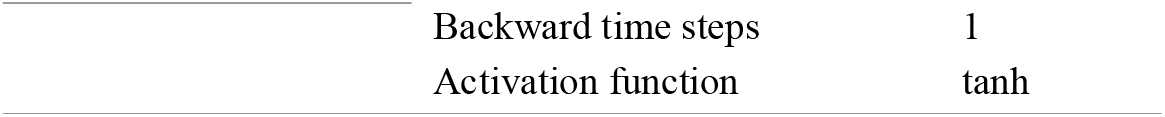
The internal structure of the learning methods used in the proposed SAAD.

**Table 2:** The classification performance of the proposed SAAD system (in terms of detection delay and ACC) using four learning methods obtained for 60-second trials and 100 repetitions of the algorithm.

| Learning Methods |  | Detection Delay | ACC |
| --- | --- | --- | --- |
| RNN | | 6231ms±443 | 89.1%±0.9 |
| QL | | 8806ms±671 | 84.1%±1.2 |
| DQL | DQL (DNN agent) | 4162ms±387 | 92.6%±0.2 |
| | DQL (RNN agent) | 2697ms±225 | 94.2%±0.4 |

Considering the dynamic nature of the learning algorithms, the system computes a detection accuracy at any time based on the cumulative decisions made up to that time. The detection delay specifies the time required for the system to reach a stable decision state in attention detection. It can be seen that the use of the QL method has the lowest accuracy and the longest delay in detection among different learning methods. Other methods using neural networks attain higher accuracies. Specifically, the use of RNN as the agent in DQL yields the highest accuracy and the shortest delay in detection. This can be interpreted by the observation that employing a powerful agent such as RNN in the internal structure of the DQL method results in higher performance of the system in terms of accuracy and detection delay.

To examine whether the high performances of the proposed dynamic SAAD system are due to the overfitting or not, the method of permutation test is used. For this purpose, the attentional labels are randomly permuted and the learning operations are repeated with the manipulated data. The results of this test for 100 repetitions are shown in Figure 5 as normal distributions for different learning approaches.

**Figure 5:**
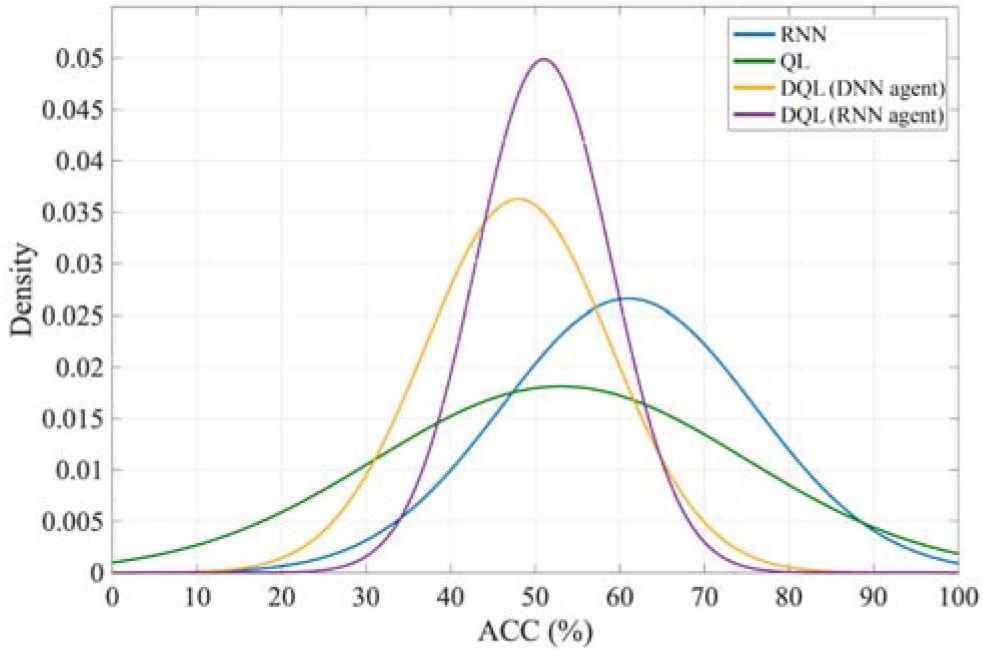
The results of the permutation test for different learning methods obtained for 100 repetitions of the proposed SAAD algorithm.

**Figure 6:**
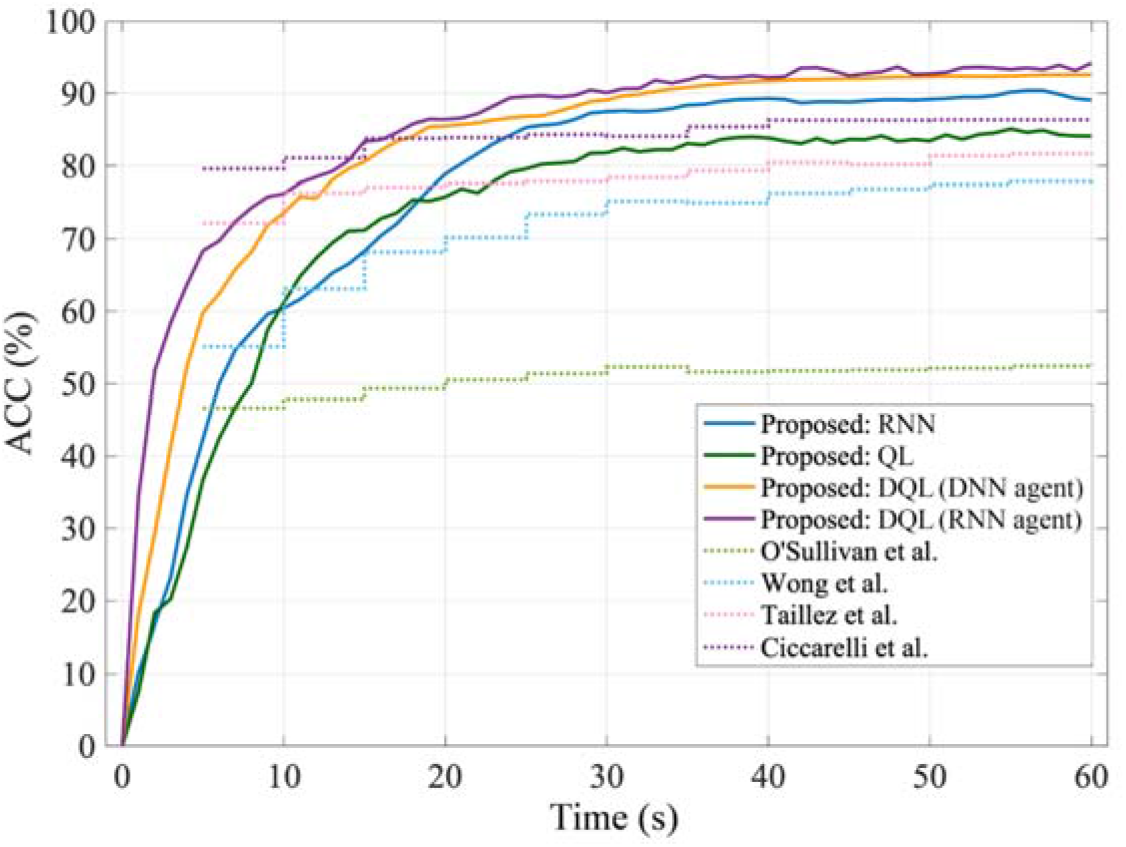
The comparison of the proposed SAAD and baseline methods in terms of ACC for sequentially increasing data duration.

According to this figure, it can be seen that using the relabeled data in the RNN method, the average detection accuracy is above 60%, which means that the overfitting in this learning method is more probable among other methods. In the QL method, despite the fact that the average accuracy is close to the level of chance, a higher variance is observed as compared with RNN for the accuracy of the detection. This indicates that these dynamic learning methods (i.e., RNN and QL) are less generalizable. However, in the deep Q-learning methods, the close values of average accuracies to the chance level (47.2% for DQL (DNN agent), 51.8% for DQL (RNN agent)) and low values of the variances, show that these methods are robust against overfitting, and therefore, suitable candidates for the dynamic SAAD system.

The second experiment concerns the comparisons of the proposed dynamic SAAD system with several traditional baseline methods (see Sec. 3.1) in terms of classification accuracy. In the first step, Table 3 shows the performances of these baselines reported in the literature with different signal lengths, along with those of the proposed system simulated with a 60-second signal length. Nevertheless, a meaningful and valid comparison is accomplished when the proposed and baseline methods are simulated under similar conditions, i.e., the same data and signal length. **Error! Reference source not found.** shows the simulation results of the methods for sequentially increasing data duration. In this diagram, for the static baseline methods, the signal length used to find the accuracy of the detection is increased in 5-seconds steps. In the case of the proposed dynamic system with different learning methods, the signal length is increased continually, where each point on the diagram represents the ratio of total true decisions to all decisions obtained up to that point.

**Table 3:** The comparison of the proposed SAAD and baseline methods (in terms of ACC). The classification accuracies of the baselines are reported from the corresponding literature.

| SAAD Method | Learning Method | Signal | ACC |
| --- | --- | --- | --- |
| O'Sullivan et al. | Reconstruction | 30 min | 89.0% [10] |
| Wong et al. | Regularized TRF | 30 sec | 90.9% [27] |
| Taillez et al. | DNN | 60 sec | 97.6% [33] |
| Ciccarelli et al. | CNN | 10 sec | 87.0% [34] |
|  | RNN | 60 sec | 89.1% |
| Proposed | QL | 60 sec | 84.1% |
|  | DQL (DNN agent) | 60 sec | 92.6% |
|  | DQL (RNN agent) | 60 sec | **94.2%** |

Evidently, the method of “Ciccarelli et al.” achieves the highest performance, whereas “O’Sullivan et al.” performs the least among the different static methods. In contrast to the result presented by “O’Sullivan et al.” in Table 3, here the method yields a final lower classification accuracy (~52%) due to the small amount of data (i.e., 60 s) used by the method. Despite the observation that the dynamic learning methods yield stable detection accuracies with some delays, the corresponding values of accuracies are, in general, higher than those of static baseline methods. Specifically, the proposed SAAD based on the DQL (RNN agent) learning method attains the highest accuracy among all dynamic SAAD approaches. The proposed system, in this case, has also the fastest rate of increase in accuracy regarding its lower detection delay (refer to Table 2). Moreover, the diagram illustrates that both the DNN and RNN agents in DQL result in close performances of attention detection in longer durations of data.

In the last experiment, considering the capability of the proposed dynamic SAAD system in temporally tracking the auditory attention, the performance of the system in switching attention scenarios is evaluated. Due to the high efficiency of DQL (RNN agent) in the previous experiments, this learning method is used to inspect the capability of the proposed SAAD for the switching attention task. The results of this experiment are shown in Table 4 for different time segments of the data (see Sec. 2.2.1). The evaluation is performed in terms of the classification accuracy of the model and the detection delays of the learning method for different time fragments of input data. As can be seen, the length of data pieces and detection accuracy are directly related to each other; the smaller the duration of the data segments, the lower the accuracy of the detection. This observation is justified by the evidence that the reduction in the length of data segments results in a decrease in classification accuracy, which is also confirmed by the findings given in **Error! Reference source not found.**. Assuming the approximately constant detection delay of the learning method (~2.5 s), the results of the table also show that even with short data segments (e.g., 5 s), an acceptable detection accuracy (~59.7%) is achievable.

**Table 4:** The classification performance of the proposed SAAD system with DQL (RNN agent) (in terms of detection delay and ACC) in attention switching scenarios obtained for different data segments and 100 repetitions of the algorithm.

|  | Data Segments |  |  |  |
| --- | --- | --- | --- | --- |
| | 12 × 5 s | 6 × 10 s | 4 × 15 s | 2 × 30 s |
| **ACC** | 59.7% ±0.3 | 73.4% ±0.2 | 84.6 ±0.2 | 90.8 ±0.1 |
| **Detection Delay** | 2432ms±216 | 2486ms±212 | 2531ms±218 | 2713ms±214 |

## 4. Conclusion

In this paper, a new system for selective auditory attention detection based on dynamic learning methods is proposed with the ability of temporally tracking the attentional direction of the listener. The main contribution of this study is to formulate the classification problem of selective auditory attention as a sequential decision-making process which is solved by the learning methods of RNN, QL, and DQL. In the proposed SAAD system, first, a set of all possible states corresponding to the input data (i.e., speech and EEG), called the probabilistic state space, is formed at each time step. Then, the generated states at each moment are used in the learning stage to detect the attentional direction. Here, various dynamic learning methods, including RNN, QL, and DQL are employed to make the final decision of the attention detection task. The proposed model for attention detection is advantageous in some aspects to the traditional auditory attention detection procedures. First, the auditory attention is detected at any time for the sequentially presented input data without requiring a separate training stage of the classifier. This means that the learning process takes place in little time, specified by the detection delay, using trial and error methods. Furthermore, the proposed SAAD model provides the possibility to be employed in such environmental conditions where the switching attention of the listener takes place. Due to the sequential and real-time nature of the learning methods in the new system, here, both the computational load and the data size are significantly low.

Using different learning methods, the proposed SAAD model is evaluated and compared with different traditional attention detection methods from the literature used as baselines. The permutation test is used to validate the reliability of the classification results and the generalizability of the methods. As a result of the experiments, it is found that the deep *Q*-learning method using RNN as agent (i.e., DQL (RNN agent)) has the best performance in terms of all criteria, including highest classification accuracy (~94.2%), least detection delay (~2697 ms), and better generalizability among all learning methods. In an additional experiment, taking the capability of the proposed SAAD in tracking the auditory attention over time, the performance of the system is assessed in switching attention conditions. The results of using DQL (RNN agent) as the learning method demonstrate that the SAAD model is able to track the switching attention of the listener for different time segments of the data. Specifically, for short data segments, the model shows a good performance in tracking the switching attention, which increases by using longer segments of data.

In a real-world cocktail party scenario, the selective auditory attention detection can be considered as a complementary component of a complete speech segregation system for the design of the hearing aid devices. The current work evaluates the performance of the dynamic SAAD system in a dichotic scenario with two input speech signals. Also, artificially produced speech segments are used to evaluate SAAD in attention switching conditions. The assessment of the proposed system with more sound sources located at different spatial positions in a noisy and reverberant acoustic environment seems to be an indispensable step toward the design of practical hearing aids. In future works, more realistic data are required to exploit the capability of the proposed dynamic SAAD system in detecting and tracking the auditory attention switching in such real environments.

